# Ancestral reconstruction of polyethylene terephthalate degrading cutinases reveals a rugged and unexplored sequence-fitness landscape

**DOI:** 10.1101/2024.04.25.591214

**Authors:** Vanessa Vongsouthi, Rosemary Georgelin, Dana Matthews, Jake Saunders, Brendon M. Lee, Jennifer Ton, Adam M. Damry, Rebecca L. Frkic, Matthew A. Spence, Colin J. Jackson

**Author notes:** These authors contributed equally to this work (Vanessa Vongsouthi, Rosemary Georgelin, Dana Matthews).

## Abstract

The enzymatic degradation of polyethylene terephthalate (PET) is a promising method of advanced plastic recycling. Traditional protein engineering methods often fall short in exploring protein sequence space for optimal enzymes due to structural and rational design limitations. Our study addresses this by using multiplexed ancestral sequence reconstruction (mASR) to explore the evolutionary sequence space of PET-degrading cutinases. With a dataset of 397 cutinase sequences, we created a diverse library of ancestral sequences. Experimental characterization of 48 ancestral sequences revealed a wide range of PETase activities, highlighting the value of mASR in uncovering functional variants when compared to traditional ASR. Our results showed that PETase activity in cutinases evolved through diverse pathways involving distal mutations to the active site, and is readily accessible within this family. Additionally, our analysis of the PETase fitness landscape using one-hot encoding (OHE) and local ancestral sequence embedding (LASE) highlighted the effectiveness of LASE in capturing sequence features relevant to activity. This work emphasizes the utility of mASR as a protein engineering tool for identifying enhanced PET-degrading enzymes, and the advantages of the LASE embedding scheme in mapping the PETase fitness landscape.

## Introduction

Over 350 million tonnes of plastic are produced annually, with the majority being derived from petrochemicals^1^. However, the current global recycling rate of plastics is estimated to be less than 10%, partially due to the technological constraints of mechanical (melt-extrusion) recycling^2^. There is a pressing need to develop advanced and scalable recycling methods that will enable the transition towards a circular plastic economy. In the last decade, enzymatic depolymerization of plastics has emerged as one such advanced recycling method^3^. The enzymatic degradation of polyethylene terephthalate (PET), a versatile thermoplastic commonly found in food and beverage packaging, and polyester textiles has received particular interest^3^.

PET-degrading enzymes that have been characterized to date, including those classified as cutinases^4–6^ (EC 3.1.1.74), lipases^7^ (EC 3.1.1.3), and carboxylesterases^8^ (EC 3.1.1.1), belong to the esterase subclass (EC 3.1) and are characterized by a catalytic triad (Ser-His-Asp) typical of the α/β hydrolase fold superfamily. Cutinases have garnered significant interest due to their ability to hydrolyse both aromatic and aliphatic polyesters^9^. Cutinases of bacterial^5,9,10^, fungal^7^, and metagenomic^11–13^ origins have been studied for their PETase activity, including Thc_Cut1 and Thc_Cut from *Thermobifida cellulosilytica*^5^, TfCut2 from *Thermobifida fusca*^5^, HiC from *Humicola insolens*^14^, FsC from *Fusarium solani pisi*^15^, and LCC from leaf-branch compost metagenome^16^. However, extant cutinases are often unsuitable for direct use in industrial processes, necessitating optimisation to improve properties such as catalytic efficiency^17^, stability under harsh conditions^18,19^, and to alleviate product inhibition^20^. For example, active site optimisation through rational design^6,10,21^ and evolutionarily-guided engineering^20^ has yielded significant improvements in PET hydrolysis across various cutinase backgrounds. While structure-guided protein engineering has been effective in improving the efficiency of extant PETases, it has provided little insight on the mechanisms by which PETase activity emerges or has been optimized by evolution.

Ancestral sequence reconstruction (ASR), which utilizes a phylogenetic tree and a statistical model of evolution to infer the sequences of extinct, ancestral proteins, can provide critical insights into molecular evolution and functional diversification within protein families^22^. ASR has been useful in studying sequence-function relationships over protein families^23,24^; such insight can shed light on the topologies of fitness landscapes that dictate the adaptive potential of proteins^25^. For example, ASR can give insight into the ruggedness (i.e. complexity) of a fitness landscape over large spans of evolutionarily accessible sequence space to reveal fitness peaks that are inaccessible through stepwise mutational approaches^24^. In addition to this, ASR is also frequently used to engineer enzymes with enhanced industrial properties^22,26–28^, making it a valuable tool in both protein engineering and understanding the mechanisms of molecular evolution.

More recently, protein representation learning has become a widely used method in protein engineering and evolutionary inference^29,30^. Protein language models (PLM), which are deep neural networks trained to predict the identities of masked residues in a corpus of protein sequences^29–32^, can map information sparse and high dimensional protein sequences to fixed-length vector representations. These vector representations capture the evolutionary and biophysical features of protein sequences in the representation model’s latent embedding space^29,30^. Indeed, PLMs have been used to learn the structure of fitness landscapes^33–35^, and can learn evolutionary features when trained with ancestrally reconstructed sequence data^36^.

In this study, we applied ASR to explore the evolutionarily accessible sequence space of PET-degrading cutinases. Using a dataset of 397 extant cutinase sequences with significant homology to known cutinases with PETase activity, we adopted a multiplexed ASR (mASR) approach to generate a diverse library of ancestral cutinase sequences. Through experimental characterization of 48 ancestral cutinases, we identified a broad range of PETase activities, including between equivalent nodes on distinct yet statistically indifferent phylogenetic topologies. Such findings highlight the importance of sampling diverse phylogenetic backgrounds to uncover functional ancestral variants. Furthermore, our study analyzed the topology of the PETase fitness landscape through two sequence embedding schemes: one-hot encoding (OHE) and the more recently described local ancestral sequence embedding (LASE)^36^. We found that LASE was more effective in capturing cutinase sequence features pertinent to PETase activity, demonstrating a clear pattern of iterative improvement in PETase functionality throughout the sequence exploration phases of our study. This comprehensive approach not only highlights the utility of mASR in uncovering novel PETase variants but also emphasizes the role of advanced embedding techniques in mapping PETase fitness landscapes.

## Results

### Multiplexed ASR yields functional PETases from diverse phylogenetic backgrounds

We used ASR to explore the PETase functional sequence space of the cutinase family. To maximize diversity in the local sequence space around known PETases, such as *Tf*Cut2 and LCC, we employed the recently described multiplexed ASR (mASR)^36^. In brief, mASR samples multiple statistically indifferent phylogenetic backgrounds from which to reconstruct ancestral sequences from. This produces diverse libraries of ancestral proteins that span functional sequence space over a distribution of realistic phylogenies. To achieve this, we performed 20 replicates of maximum likelihood (ML) phylogenetic inference and ASR on a single dataset of 397 extant cutinases with significant homology to *Tf*cut2 and LCC (E-value <= 1E-5). Consistent with previous phylogenetic studies^37^, PET hydrolytic cutinases were resolved as a polyphyletic group of two monophyletic lineages: the *Thermobifida sp*. PETases, which include *Tf*cut2, *T. cellulosytica* cutinase and *T. alba* esterase 1 and the LCC-like PETases that include LCC and BhrPETase. This topology was resolved consistently over all 20 phylogenetic priors used for mASR (**Figure 1A**). The placement of the PETase clades were supported by high ultra-fast bootstrap approximations^38^ (>= 0.95) and the ancestral nodes separating these groups were reconstructed in CodeML from the PAML^39^ suite with relatively high mean posterior probability (>0.9; based on ASR statistical benchmarking studies^40^). The resulting ancestral sequence library comprised approximately 1600 unique sequences that encompassed the evolutionarily accessible sequence space of the bacterial cutinase family.

**Figure 1.**
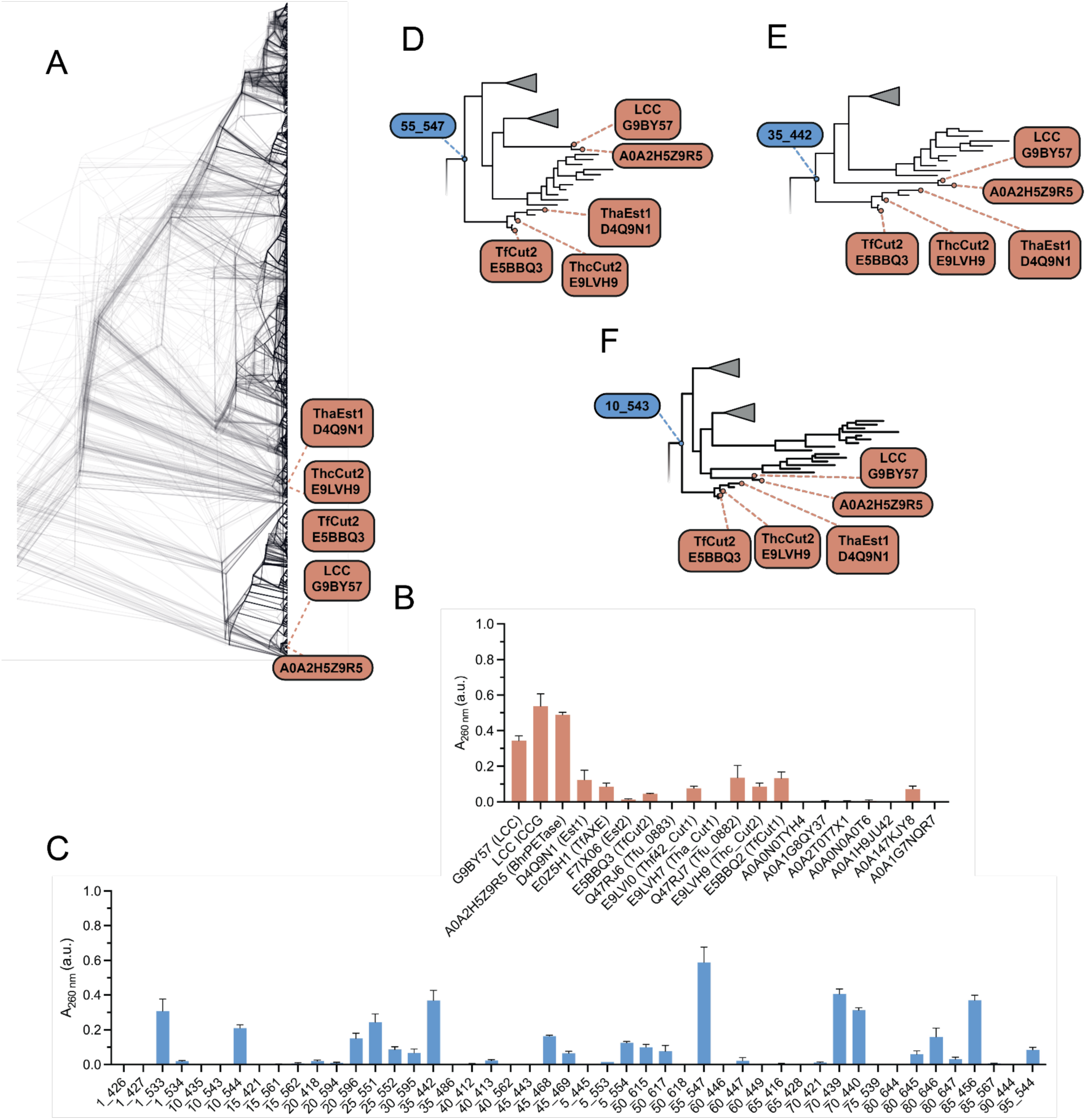
Ancestral sequence reconstruction and characterization of ancestral cutinases. (**A**) 20 replicates of phylogenetic reconstruction of the cutinase family with fixed extant node locations. All 20 presented topologies failed rejection by the AU-test and are equally valid representations of the underlying sequence alignment. Extant tips of interest (*Thermobifida* cutinases, E5BBQ3, D4Q9N1, E9LVH9 and metagenomic assembled cutinases A0A2H5ZR95, G9B757 (LCC)) are labeled). Degradation of amorphous PET film by. (**B**) 20 extant and (**C**) 48 ancestral cutinases. The bulk soluble products of PET hydrolysis (TPA, MHET, BHET) are measured by absorbance at 260 nm after 16 hours incubation with the enzyme at 60 °C. Data are represented as the mean ± SEM (n = 3). (**D**) Phylogenetic tree 55, (**E)** 35 and (**F**) 10, with ancestors 35_442, 55_547 and 10_543 highlighted. Each ancestor belongs to equivalent nodes from independent phylogenetic topologies, and are the most recent common ancestor of LCC and TfCut2, extant cutinases with known PETase activity.

We selected 48 ancestral nodes from 20 different trees with homology to the most recent common ancestor of LCC and *Tf*Cut2 for experimental characterization and comparison to extant cutinases. Specifically, these nodes were chosen from the recent ancestors of the LCC and *Tf*Cut2 lineages (or the most recent common ancestor of both), with at least a single sequence sampled from each of the 20 distinct phylogenetic trees. Each ancestral sequence was selected as the *maximum a posteriori* (MAP) sequence that maximizes the posterior probability over the full-length of the reconstructed protein. The variants were expressed, purified and tested for PETase activity. The soluble expression level of each variant was measured using the Bradford assay (**Supplementary Figure 1**). PETase activity against amorphous PET film was measured by UV absorbance at 260 nm to detect the soluble products of PET hydrolysis after the removal of undigested film, including terephthalic acid (TPA), mono(2-hydroxyethyl) terephthalate (MHET), and bis(2-hydroxyethyl) terephthalate (BHET) (**Supplementary Figure 2**). Among the extant cutinases, the highest PETase activities were observed for LCC (A_260_ nm = 0.34 ± 0.03), BhrPETase from bacterium HR29 (A_260_ nm = 0.49 ± 0.01), and a previously engineered variant of LCC with the mutations F243I/D238C/S283C/Y127G (LCC ICCG)^6^ (A_260_ nm = 0.54 ± 0.07), with other cutinases showing lower activity (**Figure 1B**). Of the 48 ancestral variants, we observed PETase activity for a number of ancestral cutinases from a diverse range of trees (**Figure 1C**). Several variants (1_533, 35_442, 55_547, 70_439, 70_440, and 85_456) exhibited similar activity to the most active extant cutinases, LCC and BhrPETase. From this group, we selected ancestors from tree 35, node 442 (35_442; A_260_ nm = 0.37 ± 0.06) and tree 55, node 547 (55_547; A_260_ nm = 0.59 ± 0.09), for further investigation. Notably, these two ancestors belonged to equivalent positions from two independent phylogenetic trees (Figure **1D-F**), representing the most recent common ancestor of LCC and TfCut2, and sharing 98.5% sequence identity to one another.

Within the dataset of 48 ancestral sequences, 12 represent the same phylogenetic node (the most recent common ancestor of LCC and *Tf*Cut2) over 12 different phylogenetic backgrounds. Of these, 9 exhibited PETase activity while 3 were inactive on PET. Interestingly, the PETase ancestor 55_547 and the inactive ancestor 10_543 both represent equivalent positions (the most recent common ancestor of *Tf*Cut2 and LCC) in their respective phylogenetic backgrounds and differ by only 9/261 positions, yet 55_547 has activity against PET and 10_543 does not. The functional differences between these sequences arise solely from the topology of the phylogenetic tree used to reconstruct them and drastically alters how the evolution of PETase activity could be inferred. For example, in the case of the 55_547 phylogeny, PETase activity appears to be a promiscuous ancestral trait that existed prior to the discovery of extant PETases (such as LCC and *Thermobifida* cutinases), whereas the 10_543 phylogeny supports the contradictory hypothesis that PETase activity emerged independently in LCC-like and *Thermobifida* PETase lineages from an ancestor without PETase activity. As both topologies failed rejection by the approximately unbiased (AU)^41^ test and are statistically indifferent at representing the observed alignment data, it is not possible to reject one of these hypotheses purely from a phylogenetic perspective. We also observe no correlation between the mean posterior probability of an ancestor and its activity (**Supplementary Figure 5**). This observation is somewhat counterintuitive, as the mean posterior probability of a sequence, which is the statistical confidence in the identity of a reconstructed ancestor, is often used to discriminate between poorly reconstructed sequences (and hence likely to be less fit) and those that are likely to be functional and fit^40^. Together, these results highlight the importance of sampling diverse phylogenetic backgrounds during ASR when accessing the functional sequence space of a protein family as minor phylogenetic incongruencies can translate into significant functional differences in (equivalent) reconstructed ancestral sequences.

### Structural characterization and analysis of ancestral cutinases

A comparative structural and sequence analysis of inactive and active ancestors suggests that PET hydrolysis appears to be a trait that is evolutionarily accessible from within the cutinase background. Unlike the *I. sakaiensis* PETase, which emerged through conformational optimization of the first- and second-shells *via* obvious selection within the active site^42–44^, PETase activity in the TfCut2 and LCC PETase lineages appears to emerge through non-specific and likely neutral mutations that are often distal to the active site (**Figure 2A-C**). Indeed, analysis of ancestral cutinase sequences reveals virtually no differences in the PET binding sites between PETase active and inactive variants (**Supplementary Figure 6**). Furthermore, mutations associated with a gain-of-function vary between the different phylogenetic trees used to reconstruct the ancestral sequences. For example, ancestors 55_547 and 35_442 are each separated by 9 unique mutations from their closest relatives without PETase activity (ancestors 10_543 with A15S, A36V, T49S, T92S, N109D, R114S, N145R, I180V, S226A and 75_539 with A36V, T49D, T92S, M105Q, R114S, S124N, R167T, P199S, A226S, respectively). Of these loss of function mutations, 3 are shared between either background, 5 are unique and one is a reversion (S226A for 55_547, A226S for 35_442). Intuitively, a combination (with at least one) of these mutations are required to be fixed in either respective cutinase background to impart PETase activity. Similarly, ancestor 10_543 is separated by 8 mutations from its closest relative with PETase activity (70_439 with E28Q, S49D, S124N, R145N, V180I, P199S, S215T, A226S); of the 8 mutations associated with gain-of-function in the background of 10_543, only 3 are specifically shared with gain-of-function mutations in 55_547. Nearly all functionally consequential mutations across backgrounds are distal to the active site and PET binding pocket. This suggests that (i) there are numerous (and diverse) molecular mechanisms by which PETase activity can emerge from an ancestral cutinase without PETase activity, (ii) these mechanisms are not obviously associated with a restructuring of the enzyme active site and (iii) PETase activity is readily evolutionarily accessible within the cutinase family, consistent with recent observations of PETases that have emerged from cutinase backgrounds^11,12^.

**Figure 2.**
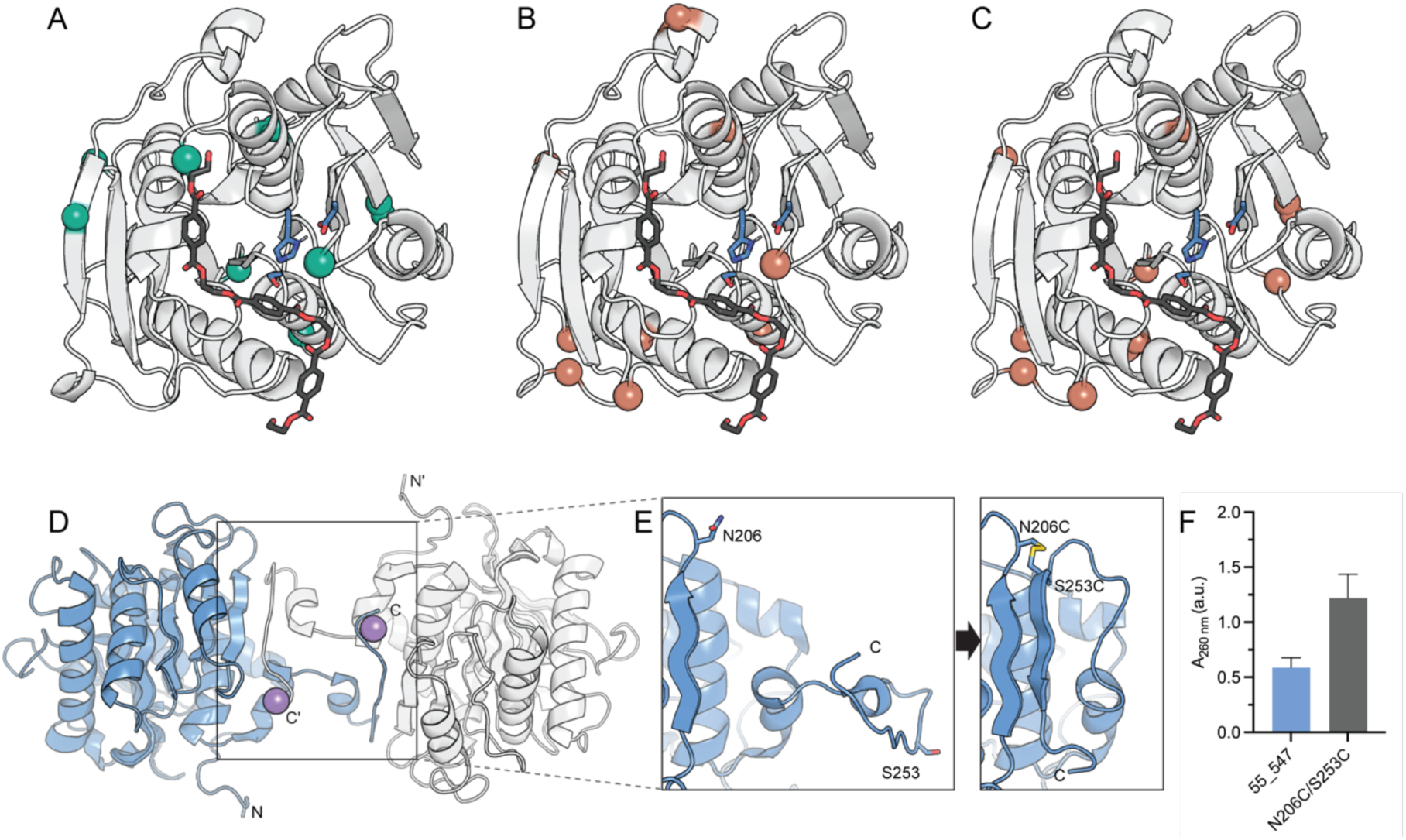
Structural characterization of 55_547 and 35_442. **A)** Positions of gain-of-function (GOF) mutations in the inactive ancestor 10_543, and **B)** loss-of-function (LOF) mutations in the active ancestors 55_547 and **C)** 35_442. GOF and LOF mutations are shown as teal and red spheres, respectively. AlphaFold^45,46^ was used to generate models of each ancestor. Most GOF/LOF mutations are distal from the catalytic site residues (blue), and the putative PET binding site highlighted by the docked 2HE-(MHET)3 ligand (black). Structural analysis of the active site residues H210, S132 and D178 in ancestors 10_543, 55_547, and 35_442 shows uniform alignment, indicating that PETase activity does not arise from modifications within the active sites (**SI. Figure 4**). **D)** Domain-swapped homodimer observed in the crystal structure of ancestor 55_547 with two C-terminal strands swapped (PDB 8ETX). The N- and C-termini of each subunit are indicated. The dimer is coordinated by sodium ions (represented as purple spheres) in the crystal lattice. **E)** Positions on the flexible C-terminus of ancestor 55_547 targeted for disulfide engineering, as shown in the crystal structure (left; PDB 8ETX), resulting in the disulfide mutant N206C/S253C, as generated by AlphaFold^45,46^ (right). The predicted formation of an additional β-strand at the C-terminus, constrained by the disulfide bond, is shown. **F)** PETase activity of ancestor 55_547 and the disulfide mutant N206C/S253C. The bulk soluble products of PET hydrolysis (TPA, MHET, BHET) are measured by absorbance at 260 nm after 16 hours incubation with the enzyme at 60 °C. Data are represented as the mean ± SEM (n = 3).

We next determined the crystal structures of two active ancestors, 35_442 (PDB 8ETY) and 55_547 (PDB 8ETX), and identified an additional pathway to optimizing PETase activity beyond modification of the active site. Both ancestors crystallized in the C222 space group at resolutions of 1.5 - 1.8 Å (**Supplementary Table 1**). Structural analysis revealed the formation of domain-swapped dimers (**Figure 2D**). The observation of domain swapping, likely induced by the high protein concentration in the crystallization conditions, suggests an intrinsic flexibility of the C-terminus, a characteristic commonly observed in proteins previously characterized to form domain-swapped dimers^47^. Given the flexible nature of the C-terminus, it was hypothesized that introducing a disulfide bond to constrain this region into forming an intramolecular β-sheet would enhance enzyme stability (**Figure 2E**). Furthermore, the chosen site for the disulfide bond coincided with the predicted Ca^2+^/Mg^2+^ binding site of the ancestors based on homology to extant cutinases, a region previously targeted for disulfide bond engineering in TfCut2 and LCC for improved thermostability and PETase activity^6,48,49^. We introduced the disulfide mutation N206C/S253C to ancestor 55_547 and experimental characterization of the resulting mutant demonstrated an approximately 2-fold improvement in whole-cell activity (A_260 nm_ = 1.22 ± 0.22) (**Figure 2F**) and a 1.5-fold increase in soluble expression levels compared to the 55_547 background determined by Bradford assay (**Supplementary Figure 5**). In combination with our comparative analysis of the selected inactive and active ancestors, the successful optimization of PETase activity via the introduction of a disulfide bond distal to the active site illustrates that PETase activity within the cutinase family can emerge and be enhanced through mechanisms that extend beyond the structural reconfiguration of the active site. It is therefore likely that activity optimization is, at least in part, being driven by thermodynamic and kinetic stabilization.

### Alternate Reconstructions of Ancestral Cutinases 35_442 and 55_547

We next experimentally characterized all alternate sequences of ancestors 55_547 (40 mutants) and 35_442 (29 mutants) that individually sampled each mutation that had been ambiguously reconstructed. Here, we define ambiguity as any site where at least two amino acids are reconstructed with a posterior probability of >= 0.2. This provided a high resolution mutagenic map of the local sequence space around either of the ancestral PETase variants. Ambiguously reconstructed sites were spatially distributed over the protein and not localized to any specific functional area (**Supplementary Figure 7**). Having already established that the mean posterior probability of an ancestral sequence is a poor indicator of PETase activity (**Supplementary Figure 4B**) and that relatively minor changes in the cutinase sequence can drastically alter PETase activity, we hypothesized that the neighborhood of evolutionarily possible (albeit less probable) sequences may contain mutations that benefit PETase activity. This hypothesis was guided by recent ASR studies on the *I. sakaiensis* PETase branch of the cutinase phylogeny where PETase activity appeared to emerge transiently within ancestral lineages^50^. Experimental characterization revealed that all of the alternate reconstructions of ancestors 55_547 and 35_442 demonstrated PETase activity that was comparable (or reduced) to either respective ancestral background (**Figure 3A-B**).

**Figure 3.**
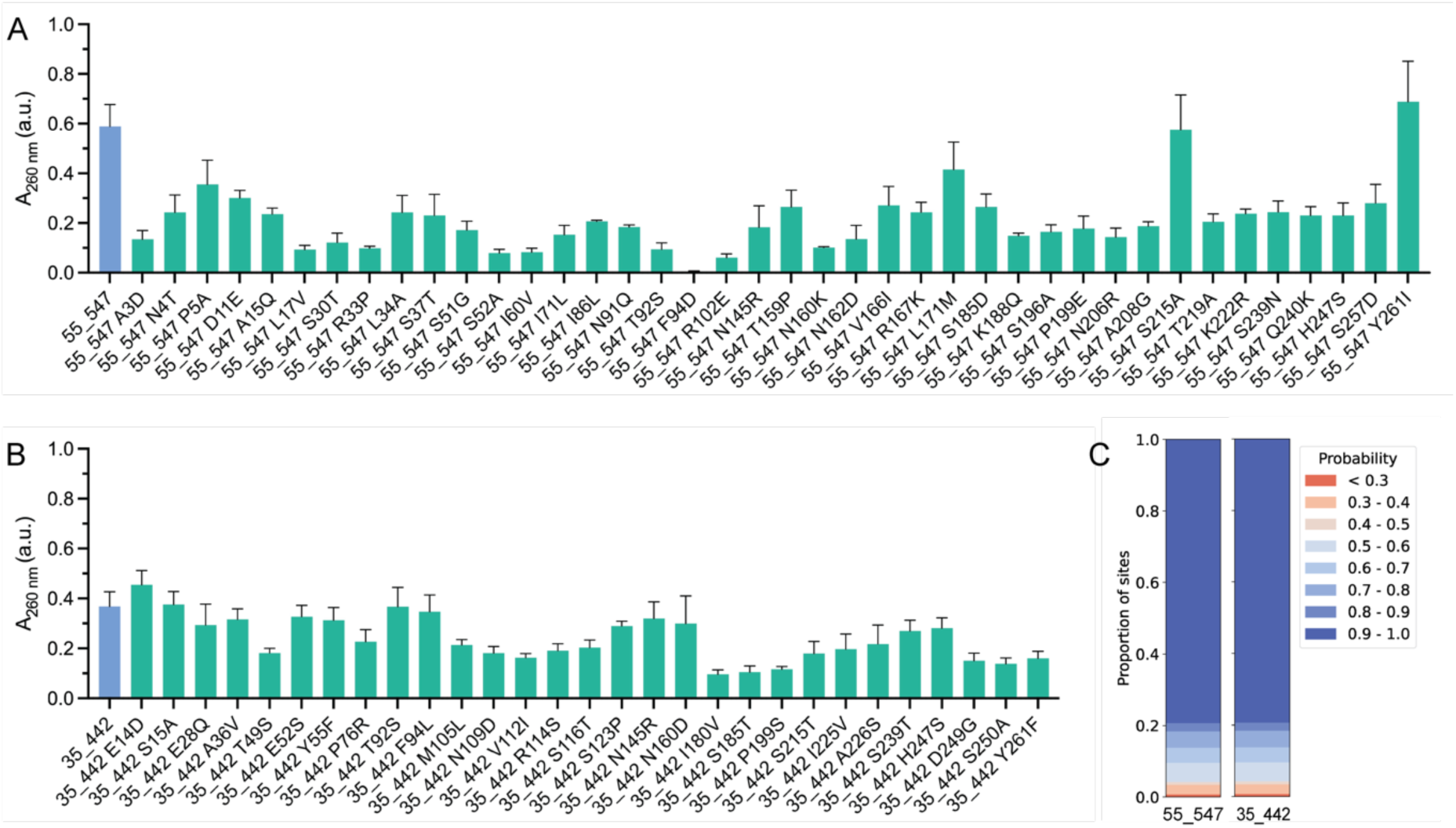
Alternate reconstructions and random recombinations of ancestral cutinases 35_442 and 55_547. Activity of alternate reconstructions of **A)** ancestor 55_547 and **B)** 35_442 against amorphous PET film. The bulk soluble products of PET hydrolysis (TPA, MHET, BHET) are measured by absorbance at 260 nm after 16 hours incubation with the enzyme at 60 °C. Data are represented as the mean ± SEM (n = 3). **C)** Posterior probability distributions for ancestors 55_547 and 35_442.

A subset of randomly sampled combinations of single mutations from the alternate reconstructions were selected to explore potential epistatic interactions and their impact on PETase activity. The 7 selected mutations, E14D, E28Q, A36V, S52A (in 55_547), F94L, S196A, and H247S, were recombined in various combinations from 2 to 5 point mutations in the background of 35_442 and 55_547. E28Q, A36V, S52A, S196A, and H247S are surface mutations, distal from the catalytic and putative PET binding site (**Figure 4A**). In contrast, F94L is positioned within the PET binding site, as deduced from docking of 2HE-(MHET)_3_ into both 35_442 and 55_547 (**Figure 4A**), as well as structural homology to previously predicted PET binding sites in LCC. As single mutations, E14D, E28Q, A36V, F94L and H247S were considered neutral based on PETase activity relative to the 35_442 background, while S52A and S196A exhibited decreased activity relative to 55_547. In our experimental characterization, we identified 24 recombined variants with increased PETase activity relative to their ancestral background (**Figure 4B**). Specifically, 10 recombined variants in the background of 55_547 showed increased PETase activity, with the most active being E14D/E28Q/A36V (A_260_ nm = 1.20 ± 0.26), E28Q/S196A (A_260_ nm = 1.08 ± 0.05), and E28Q/S52A/S196A (A_260_ nm = 1.03 ± 0.09). Similarly, 14 recombinations in the background of 35_442 demonstrated increased activity, with the most active being E28Q/S196A (A_260_ nm = 1.42 ± 0.29), E28Q/F94L (A_260_ nm = 0.99 ± 0.17), and E14D/H247S (A_260_ nm = 0.77 ± 0.08). Interestingly, we identified 17 recombined variants that exhibited improved PETase activity relative to LCC ICCG, the most active extant variant in the study, with 35_442 E28Q/S196A, displaying ∼2.5-fold higher activity.

**Figure 4.**
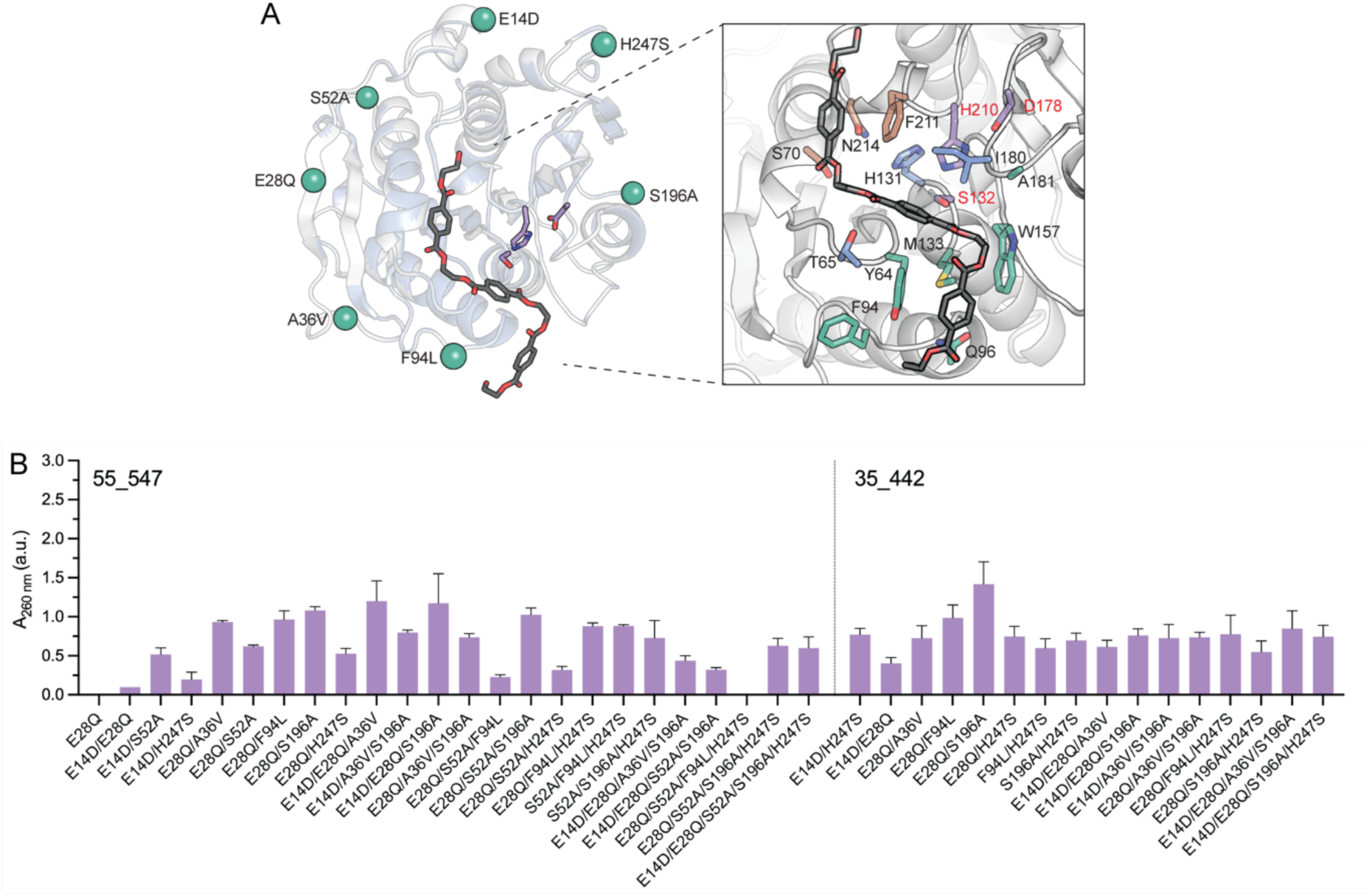
PETase activities of recombined alternate reconstructions. **A)** Structure of 35_442 and 55_547 aligned. Positions selected for recombination are highlighted as spheres. The docked pose of 2HE-(MHET)3 is shown in a close-up of the binding and catalytic site, and residues that form the putative PET binding site based on homology to LCC are shown. Residues comprising subsites -2 (green), -1 (blue) and +1 (brown) are highlighted. The catalytic residues Ser132-His210-Asp178 are also shown (purple; red labels). **B)** Activity of recombined mutations from alternate reconstructions of 35_442 and 55_547 against amorphous PET film. The bulk soluble products of PET hydrolysis (TPA, MHET, BHET) are measured by absorbance at 260 nm after 16 hours incubation with the enzyme at 60 °C. Data are represented as the mean ± SEM (n = 3) and grouped based on mutations recombined in the background of 55_547 (left) and 35_442 (middle).

The mutational analysis of the recombined variants highlighted E28Q as a key mutation, present in 14 of the 17 variants that demonstrated enhanced activity compared to the engineered LCC ICCG variant. This suggests a positive effect of E28Q on PETase activity; however, this was only observed in the presence of other mutations from the recombinations. The context dependence of E28Q is especially significant when considering the double mutation E28Q/S196A, which was among the most active in both ancestral backgrounds. When assessed independently in the 55_547 background, the E28Q mutation adversely affected activity (A_260_ nm = 0.00 ± 0.00), and the S196A mutation similarly led to a reduction in activity relative to the ancestor (A_260_ nm = 0.16 ± 0.02). However, the combination of these mutations resulted in a significant increase in activity beyond the additive effects of the single mutations (A_260_ nm = 1.08 ± 0.05), indicative of positive epistatic interactions.

### Ruggedness analysis reveals epistasis in recombined mutations

We next analyzed the topology of the PETase fitness landscape over the 196 cutinase sequences characterized in this study. This was done both to build a holistic overview of PETase evolution and function in the cutinase family and to assess the effectiveness of this approach in exploring new-to-nature fitness peaks across functional sequence space. To achieve this, we embedded cutinase sequences as nodes in a network graph where edges connect each node to its *k* nearest (Euclidean) neighbors. The scheme used to embed sequences therefore dictates the topology of the network graph. When this is the one-hot encoding (OHE), the Euclidean distance between nodes in the network graph is proportional to the number of mutations between the sequences they represent. The OHE graph network therefore captures PETase activity as a function of the mutations between sequences when the signal over the graph is the measured PETase activity (**Figure 5A**).

**Figure 5.**
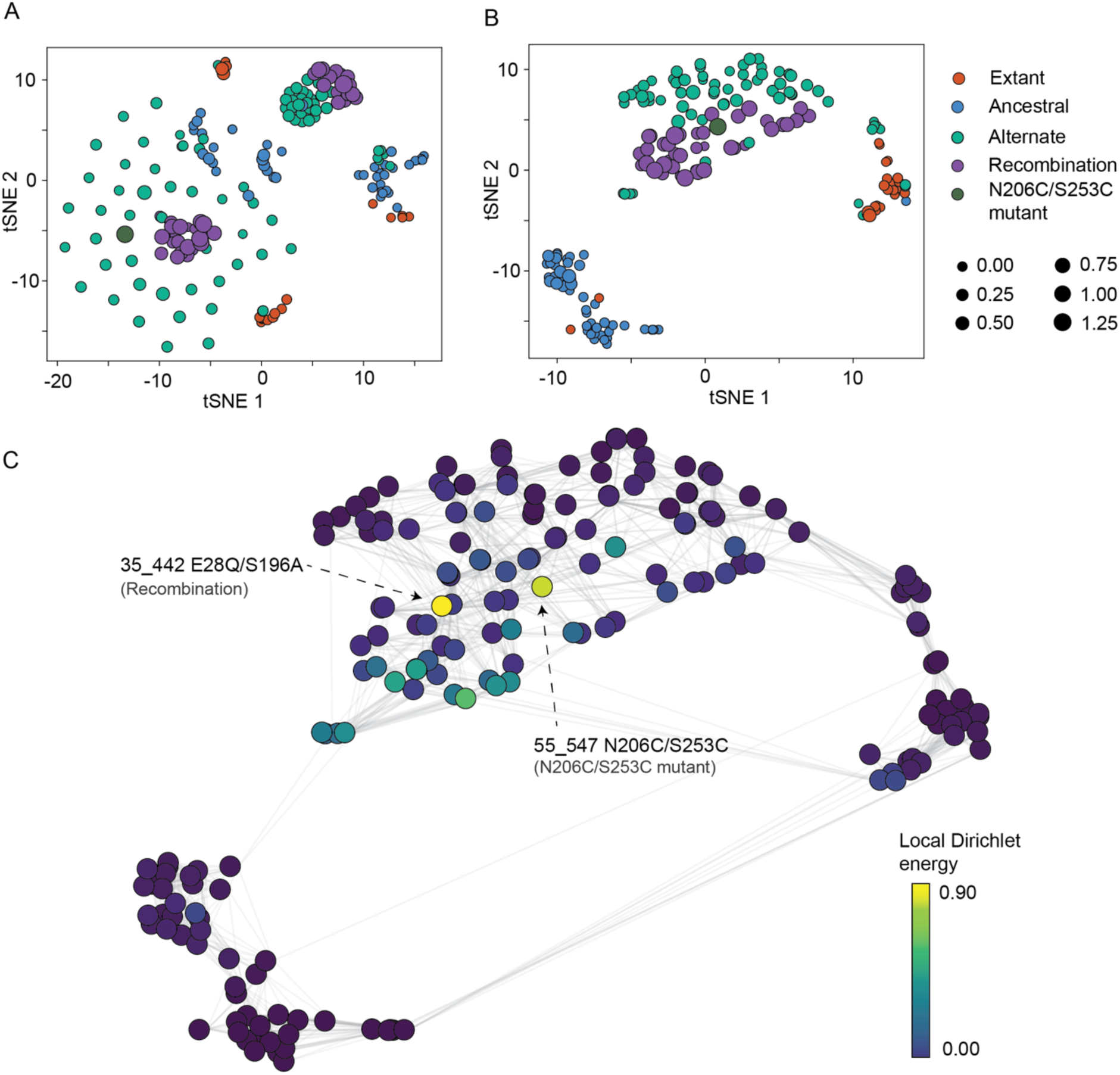
Analysis and regression on PETase sequence space. PETase sequences were represented in **A)** OHE- and **B)** LASE-forms and projected into a 2-dimensional space with tSNE. Colour represents design stage and size represents activity (A_260 nm_ (a.u.)). The OHE and LASE sequence data was then used to train regression models. **C)** The local Dirichlet energy for each variant was determined over subgraphs that include the variant’s immediate neighbors as determined by kNN. Edges connect variants that were found to be neighbors and color represents the local Dirichlet energy calculated.

Encoding protein sequences as the hidden states of a PLM can capture comparatively richer features than the OHE^29,30^, albeit at the cost of interpretability over the network graph^36^. Euclidean distances in the latent space of a representation model may share no interpretable relationship with equivalent Euclidean distances in the OHE space, therefore confounding the interpretation of PETase activity over the network graph to non-linear distances instead of simple mutational distances.

We embedded all characterized cutinases in a OHE basis and visualized the resulting landscape in 2 dimensions with t-distributed stochastic neighbor embedding (tSNE) dimensionality reduction. In this space, ancestral and extant cutinases form homogenous, random clusters (**Figure 5A**). The alternate reconstructions of ancestors 55_547 and 35_442 and their recombinations are grouped as closely connected clusters around their respective ancestral backgrounds. Despite their comparable PETase activity, sequences in both backgrounds are resolved as independent components that share few direct edge connections, suggesting that the PETase fitness landscape is multi-modal (i.e. there are multiple solution spaces to competent PETase activity) when considering only naive, residue-wise OHE sequence embeddings.

We then reconstructed the PETase fitness landscape in a learned representation space. To ensure that local features of the cutinase sequence space were captured by a PLM, we used a recently described method of local ancestral sequence embedding (LASE)^36^. In brief, LASE trains a small and family-specific deep learning model on ancestrally reconstructed sequence datasets. The features learned by LASE capture the functional properties of proteins, such as catalytic efficiency, in a non-linear space that is not interpretable with simple mutational distances. Embedding in LASE can therefore reveal topological features of the fitness map that are obfuscated in an OHE embedding.

Sequences in the LASE embedding space cluster according to their relative fitness and how they were sampled (e.g. extant, ancestral, alternate or recombined; **Figure 5B**). For example, all point mutations from alternate reconstructions and recombinations group together in a single connected component in the LASE space, irrespective of the genetic background (ancestors 55_547 or 35_442) they were introduced into. Moreso, ancestral and extant sequences cluster into disconnected components, despite their often high degree of site-wise sequence similarity. Indeed, the LASE representation space is highly structured relative to the OHE space, where ancestral and extant sequences co-occur across distinct identity groups in the network graph. This indicates that our sequence sampling strategy is highly structured and systematic over an evolutionarily informative representation space, while appearing somewhat random in the OHE space.

Finally, we used a graph signal processing approach to quantify ruggedness in the cutinase PET fitness landscape. We define ruggedness as the non-linearity between the fitness of a sequence and its neighbors in the network graph. We use the Dirichlet energy of the graph, which describes the non-linearity of a signal over a graph, to measure this^36,51,52^. In order to make the Dirichlet energy interpretable as a node-wise local quantity, we calculate it over each subgraph in the network defined by an edge-length of exactly 1, thus reducing its interpretation to the deviation from linearity a node demonstrates relative only to its immediate neighbors; the fitness signal over the network graph changes as a linear function (i.e. is smooth) over nodes of the graph that are characterized by low local DEs. The local Dirichlet energy is therefore a descriptor of how confounded by epistasis a sequence is. This analysis revealed that the fitness landscape is most rugged over the cutinase variants with the greatest PETase activity. Indeed, the node with the single highest local Dirichlet energy is also the fitness peak (ancestor 55_547_E28Q/S196A). Importantly, this was true for both OHE and LASE network graphs (**Figure 5C**; **Supplementary Figure 9**), indicating that the combinatorial mutations E28Q/S196A in the 55_547 background would be unlikely to be introduced through a stepwise mutagenesis approach due to the relatively low activity of each single mutation in isolation. Together, these analyses demonstrate that mASR can effectively guide protein engineering by discovering evolutionary features that are not immediately apparent, and can help navigate rugged regions of sequence space to find fit enzyme variants that are not rationally obvious.

## Discussion

In our investigation into the evolutionary sequence space of bacterial cutinases with PETase activity, we employed multiplexed ASR^36^ to enhance our exploration. This method allowed us to reconstruct and analyze ancestral cutinases across 20 diverse phylogenetic topologies, moving beyond the constraints of a single-tree perspective. Our experimental characterizations of 20 extant and 48 ancestral cutinases unveiled a wide spectrum of activities against amorphous PET film (A_260_ nm = 0.00 - 0.59). Of particular interest were two ancestors, 35_442 (A_260_ nm = 0.37 ± 0.06) and 55_547 (A_260_ nm = 0.59 ± 0.09), each representing the most recent common ancestor of LCC^13^ and TfCut2^5^ on independent trees and exhibiting similar activity to the most active extant cutinases, LCC (A_260_ nm = A_260_ nm = 0.34 ± 0.03) and BhrPETase (A_260_ nm = 0.49 ± 0.01). Notably, the use of mASR was crucial in uncovering the variability of ancestrally reconstructed sequences, as it revealed that equivalent nodes from independent phylogenetic topologies exhibited highly varied activity despite all topologies being equally valid representations of the underlying sequence alignment based on the AU-test. While some ancestors were inactive, others showed significant PETase activity comparable to characterized extant PETases. These observations highlight the utility of the mASR method in improving both the success and robustness of ASR applications in protein engineering. By embracing a wider array of evolutionary scenarios, mASR allowed for a more comprehensive and reliable identification of functional sequences relative to a single-tree approach. Multiplexed ASR may therefore become an important component of ASR- and evolutionary-guided enzyme design strategies in the future.

Our sequence and structural analysis of active and inactive ancestral cutinases provided insights on the evolutionary emergence of PETase activity in the cutinase family. In particular, our findings suggest that PETase activity in the TfCut2 and LCC PETase lineages has emerged through neutral mutations that are distal to the active site, rather than specific changes localized to the active site itself, as observed in the evolution of *I. sakaiensis* PETase^37^. Our detailed examination of ancestors 35_442 and 55_547, along with their closest inactive counterparts, highlighted unique gain-of-function mutations leading to PETase activity from these distinct phylogenetic backgrounds. These findings imply the existence of multiple, distinct evolutionary pathways for acquiring PETase functionality and indicate that PETase activity is readily accessible within the cutinase family.

Our structural analysis also revealed the formation of domain-swapped dimers between adjacent symmetry mates in the crystal structures of the ancestral variants 35_442 and 55_547. While possibly an artifact of the crystallization conditions, this observation suggests a certain degree of structural flexibility in the C-terminus of these ancestors even in solution. Based on this observation, we introduced a disulfide bond in the C-terminus of ancestor 55_547 that coincided with the predicted Ca^2+^/Mg^2+^ binding site, a strategy homologous to similar successful modifications in TfCut2 and LCC that were observed to improve stability and activity^6,48,49^. Indeed, the introduction of the disulfide N206C/S253C resulted in a variant of 55_547 with an approximately 2-fold improvement in whole-cell activity and 1.5-fold improvement in soluble expression relative to the background ancestor.

We deepened our exploration of the evolutionary sequence space of the cutinase family by addressing ambiguously reconstructed positions in the initial mASR. Ambiguous positions were identified based on a posterior probability threshold of ≥ 0.2 for the second most probable residue. Introducing these 69 alternate reconstructions as single mutations into 35_442 and 55_547, we observed that most mutations were either neutral or decreased PETase activity relative to the ancestral background. However, when we recombined a random subset of the alternate reconstructions, we observed several recombinations in the background of both 35_442 (E28Q/S196A) and 55_547 (E14D/E28Q/A36V) that displayed increased PETase activity, not only relative to the initial ancestors, but also to an engineered variant of LCC with enhanced activity, LCC^ICCG^ ^6^. Notably, the most active recombinations exhibited activity improvements that exceeded the additive effects predicted from their individual mutations, highlighting positive epistatic interactions that contribute to their PETase activity.

To complement our experimental investigations, we modeled the sequence-fitness landscape of all 196 cutinase sequences characterized in our study, employing a graph signal processing approach that elucidates the complex topology of the PETase fitness landscape. This methodology involved embedding cutinase sequences as nodes in a network graph, where each node was connected to its *k* nearest (Euclidean) neighbors, thereby enabling a direct comparison between two embedding schemes, OHE and LASE. While OHE provides a straightforward mutational distance metric between sequences, it often oversimplifies the nuanced relationship between sequence variation and function. In contrast, LASE, by training a family-specific deep learning model on our dataset of ancestrally reconstructed sequences, captures the functional properties of proteins in an abstract manner, revealing non-linear sequence-activity relationships that are obscured in OHE representations.

Using LASE to map the sequence-fitness landscape, we highlighted iterative improvements in PETase activity throughout the different phases of our sequence exploration of the cutinase family. This embedding not only facilitated a deeper understanding of the functional implications of sequence variation but also allowed us to identify clusters of sequences with similar functional profiles, regardless of their evolutionary background. Moreover, our analysis of the fitness landscape’s ruggedness via LASE provided new insights into the role of epistasis in PETase activity. The local Dirichlet energy calculations revealed that sequences with the highest PETase activity were associated with the greatest ruggedness, suggesting that the most functionally optimized variants emerge from complex interplays of multiple mutations rather than from linear accumulations of beneficial single mutations. This observation highlights the significance of epistatic interactions in driving the evolutionary innovation of PETase activity, illustrating that successful enzyme variants often lie in regions of the fitness landscape that are not readily accessed through single mutational steps, but can be accessed using evolutionarily-guided approaches, such as mASR.

## Materials and Methods

### Ancestral sequence reconstruction

1000 sequences were collected from the NCBI non-redundant (nr) database with blast using LCC (UniProt: G9BY57) as seed and an e-value cutoff of 1e-5. Sequence redundancy was removed to 90% ID in CD-HIT^53^. Signal peptides were deleted using SignalP4.0^54^. Alignment was performed using the GINSI protocol of MAFFT^55^. 100 replicates of independent model parameterization and tree search (default parameters) were performed using IQTREE2^56^ on the NCI GADI supercomputer. The sequence evolution model was parameterized using ModelFinder^57^, as implemented in IQTREE2. Branch supports were determined as the ultrafast bootstrap approximation^38^ calculated to 1000 replicates, as implemented in IQTREE2. The Approximately unbiased test^41^ was conducted to 10000 replicates for all ML topologies. Empirical Bayesian ASR was performed on 20 of the 100 trees that failed rejection by the AU test in CodeML^39^ using the ML replacement matrix (LG)^58^ with rates modeled as a discrete gamma parameterized with 4 rate categories.

### Small-scale protein expression and purification

Plasmids were transformed by heat shock into chemically competent BL21(DE3) *E. coli* cells and plated onto Lysogeny broth (LB) agar supplemented with 100 µg/mL kanamycin and incubated at 37 °C overnight. A single colony was used to inoculate 1.5 mL autoinduction media supplemented with 100 µg/mL kanamycin in a 2.2 mL 96-well deep well block and grown at 1050 rpm at 37 °C for 5 hours, followed by room temperature (RT; 25 °C) for 16 hours.

Cells were harvested by centrifugation at 2000 x *g* for 15 minutes at RT and resuspended in Lysis Buffer (1X BugBuster® Protein Extraction Reagent (Merck-Millipore), 20 mM Tris, 300 mM NaCl, 1 U/ml Turbonuclease (Sigma) pH 8). The cell suspension was left to incubate at RT for 20 minutes with gentle shaking. The lysate was separated from the insoluble cell debris by centrifugation at 2250 x g for 1 hour at RT.

The clarified lysate was then diluted with 100 µl of Equilibration Buffer (20 mM Tris, 300 mM NaCl pH 8) and purified by nickel-charged IMAC using a 96-well HisPurTM Ni-NTA Spin Plate (ThermoFisher Scientific) equilibrated in Equilibration Buffer, washing the sample three times with 250 µl of Wash Buffer (20 mM Tris, 300 mM NaCl, 10 mM imidazole pH 8) and eluting with 250 µl of Elution Buffer (20 mM Tris, 300 mM NaCl, 150 mM imidazole pH 8). All centrifugation steps following addition of Wash or Elution Buffer were at 1000 x *g* for 1 minute at RT. The eluate was stored at 4 °C. Bradford assay was used to quantify the soluble expression levels post-purification (**Supplementary Figure 1**).

### UV absorbance assay for PET-degrading activity

For purified protein from 96-well expression and purification, 15 µl of the eluate from the 96-well Ni-NTA purification and 285 µl of Reaction Buffer (50 mM Bicine pH 9) was added to a clear 96-well plate. For purified protein from large-scale expression and purification, 300 µl of 100 nM enzyme in Reaction Buffer was added to a clear 96-well plate. A single disk of amorphous PET (Goodfellow ES301445) with 4 mm diameter and 0.25 mm thickness was added to each well. The plate was incubated at 60 °C for 16 hours. Following incubation, 100 µl of the reaction solution was transferred to a clear UV-transparent 96-well plate and the absorbance was measured between 240 to 300 nm in 10 nm steps using the Epoch Microplate Spectrophotometer (BioTek) (**Supplementary Figure 2**). For comparison of activity of all variants, the absorbance at 260 nm was used. Assays were repeated in triplicate for each variant.

### Assay data processing

To correct for possible systematic error between replicate data points, the mean absorbance at 260 nm for each replicate was determined. Using the mean absorbance, a scaling coefficient was assigned to the replicates with the lowest and highest mean values such that the mean of the scaled absorbance values equaled the mean of the replicate with the mid-range mean value. To ensure scaling improved consistency between replicates, correlograms of the data before (**Supplementary Figure 3**) and after (**Supplementary Figure 4**) scaling were produced to confirm the monotonic (rank) correlation between replicates was preserved.

### Cloning of TEV-PETase variants for crystallography studies

Primer pairs containing the DNA sequence for the TEV cleavage site (5’-GAAAACCTGTATTTTCAAAGC-3’) were constructed, specific to each PETase variant. PCR was performed using these primers and PETase variant genes to create mutant fragments. These fragments were reassembled using Gibson Assembly^59^ and checked through Sanger sequencing to ensure the TEV cleavage site was correctly introduced.

### Large scale protein expression and purification

The TEV-PETase ancestral variants plasmids were transformed using electroporation into electrocompetent E. cloni® cells (Lucigen) and plated onto LB agar supplemented with 100 µg/mL kanamycin. The plates were incubated overnight at 37 °C. A single colony was inoculated into a 10 mL solution of LB media with 100 µg/mL kanamycin and incubated overnight at 37 °C and 180 rotations per minute (rpm). This liquid starter culture was then added to 1 L of autoinduction media^60^ (6 g Na_2_HPO_4_, 3 g KH_2_PO_4_, 20 g tryptone, 5 g yeast extract, 5 g NaCl, 10 mL of 60% (v/v) glycerol, 5 mL of 10% (w/v) glucose, and 25 mL of 8% (w/v) lactose) with 100 µg/mL kanamycin and incubated for 24 hours at room temperature and 180 rpm. The cells were separated from the media by centrifugation at 5000 × g for 15 minutes at 4 °C and resuspended in lysis buffer (400 mM NaCl, 25 mM imidazole, 1 U/mL Turbonuclease (Sigma), and 50 mM Tris-HCl pH 8.0). The resuspended cell solution was lysed using two rounds of sonication at 50% power and pulse time for 5 minutes, with 5 minutes on ice between sonication steps. Next, the sample was centrifuged at 32000 × g for 60 minutes at 4 °C, and the soluble cell solution was separated from the insoluble cell material and filtered through a 0.45 μm pore size filter. The filtered soluble cell solution was passed through an equilibrated Nickel-charged IMAC using a 5 mL HisTrap HP (GE Healthcare Life Sciences) in Lysis Buffer. The protein bound to the column was eluted using elution buffer (400 mM NaCl, 500 mM imidazole, and 50 mM Tris-HCl pH 8.0). The protein sample was buffer exchanged to TEV reaction buffer (100 mM NaCl, 0.5 mM EDTA, 1 mM DTT, 1% (v/v) glycerol, and 50 mM Tris-HCl pH 8.0) using a PD-10 desalting column and diluted to 50 mL in this buffer. A 1 mL solution containing 1 mg/ml of purified TEV protease was added and incubated at room temperature overnight. The cleaved sample was passed through an equilibrated Nickel-charged IMAC using a 5 mL HisTrap HP (GE Healthcare Life Sciences), and the flowthrough was collected. This flowthrough was concentrated using the 3 kDa Amicon® ultra 15 mL centrifugal filters and filtered through a 0.22 μm filter. Finally, the cleaved protein was purified to homogeneity using size-exclusion chromatography, and the HiLoad 26/600 Superdex 200 column (GE Healthcare Life Sciences) was equilibrated in size-exclusion buffer (150 mM NaCl, 25 mM HEPES pH 7.5).

### Protein crystallisation and structure determination

Proteins were concentrated using the 3 kDa Amicon® ultra 15 mL centrifugal filters to 15-36 mg/ml and crystallized in 20% (w/v) PEG 3350 alongside 0.2 M salt and BisTris buffer solution. Specifically, ancestor 55_547 in 0.2 M sodium/potassium tartrate, 0.1 M BisTris propane pH 7.5 and 20% (w/v) PEG 3350; and ancestor 35_442 in 0.2 M sodium malonate, 0.1 M BisTris propane pH 6.5 and 20% (w/v) PEG 3350. The X-ray diffraction data were collected on the MX2 beamline at the Australian Synchrotron^61^. The data was processed using XDS^62^, and the phase problem was resolved with molecular replacement using the PETase WT structure (PDB: 6EQE) as the search model. The ligands and solvent molecules were removed and then used as the search model part of Phaser (CCP4)^63^. The structure was refined using phenix.refine^64^ through multiple iterative steps, and rebuilt each time with Coot^65^. The structures of ancestors 55_547 and 35_442 were deposited in the protein data bank under the PDB ID of 8ETX and 8ETY, respectively.

### Protein sequence representations

To analyze PETase sequence space One-hot embeddings (OHE) and Learned Ancestral Sequence Embeddings (LASE) were made. To produce OHE, aligned PETase sequences were converted into a (20 × 267) vectors where gaps were represented as a zero vector of length 20. The LASE embedding model was implemented as a Transformer in PyTorch 2.0.1 as previously described^36^, with three encoder blocks with 2-headed multihead attention (64 dimensions) and a feed-forward fully-connected layer (128 dimensions). The LASE embedding model was trained with a masking percent of 15%, over 100 epochs with a batch size of 32 using the Adam optimizer. Loss was determined as the categorical cross entropy loss. Performance over training was assessed with perplexity and categorical accuracy (**Supplementary Figure 8**).

### Local Dirichlet energy calculations

To estimate the local ruggedness of each PETase variant, the k-nearest neighbors of each node were used to produce a KNN sub-graph for each variant with scikit-learn 1.2.1. The kNN sub-graphs were made symmetric by defining an edge in either direction to be a single edge. The dirichlet energy was calculated as previously described^52,66^:

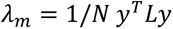

Where, *λ_m_* is normalized dirichlet energy, *N*, the number of variants in the sub-graph, *y*, the activity (absorbance) of each variant and *L* the graph Laplacian operator of the adjacency matrix of each kNN sub-graph.

## Acknowledgements

This research was undertaken in part using the MX2 beamline at the Australian Synchrotron, part of ANSTO, and made use of the Australian Cancer Research Foundation (ACRF) detector.

## Availability

The code for model training and analysis is available on GitHub: https://github.com/RSCJacksonLab/cutinase_lase

Deposited crystal structures are available on the Protein Data Bank under accession IDs 8ETY and 8ETX.

## Note

The authors declare the following competing financial interest(s): C.J.J., V.V., M.A.S., R.G., D.M., J.S., A.M.D., and J.T. hold equity in the plastic recycling company, Samsara Eco.

