## Supplementary_Information for "Ancestral reconstruction of polyethylene terephthalate degrading cutinases reveals a rugged and unexplored sequence-fitness landscape"

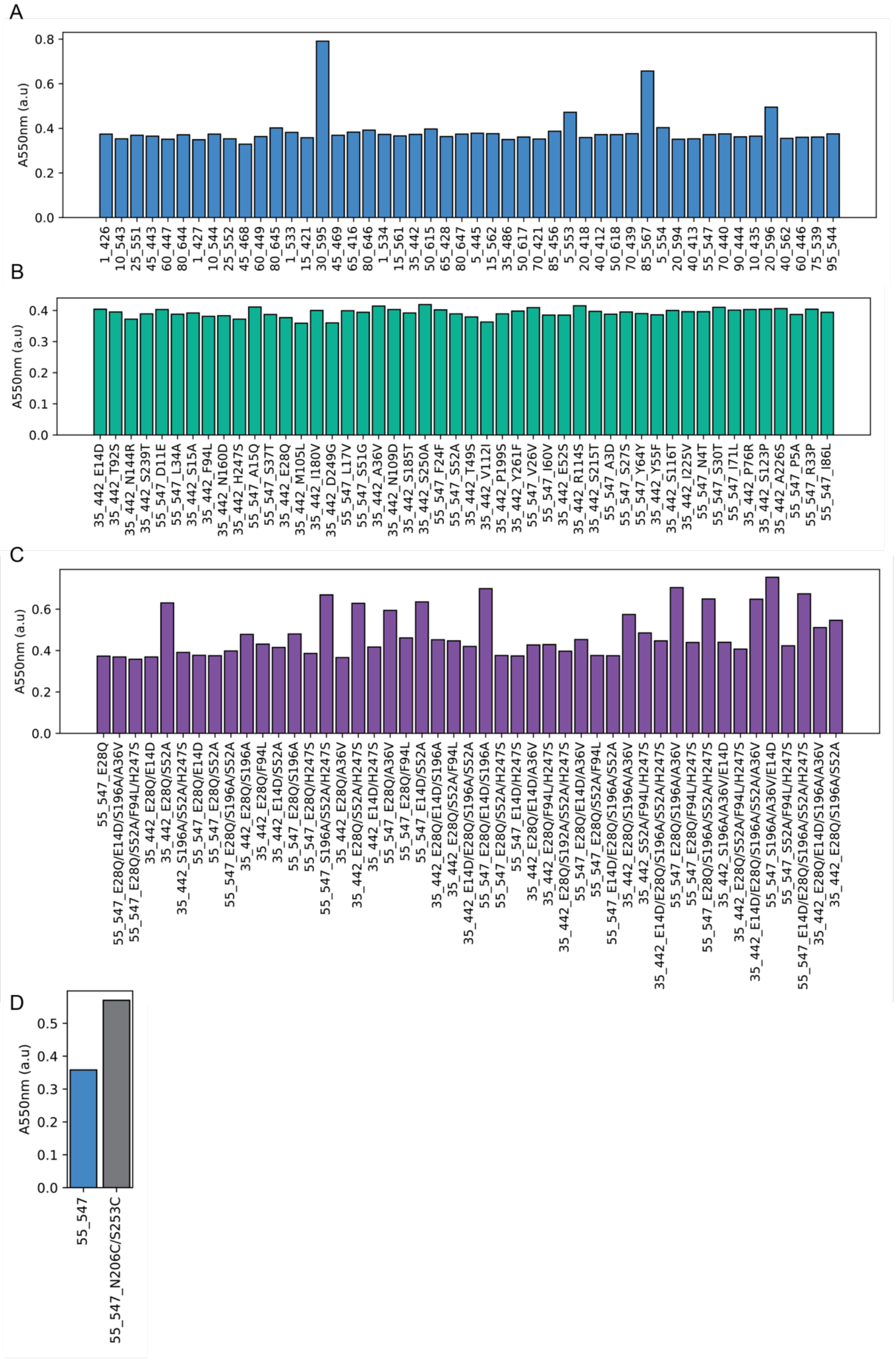


**Supplementary Figure 1.** Relative soluble protein yield post nickel immobilized metal ion affinity chromatography for **A**. ancestral cutinases, **B**. ambiguous alternate ancestors in the backgrounds of ancestors 35_442 and 55_547, **C.** recombined alternate sequences in the backgrounds of ancestors 35_442 and 55_547 and **D**. ancestor 55_547_N206C/S253C (shown with 55_547 for clarity).

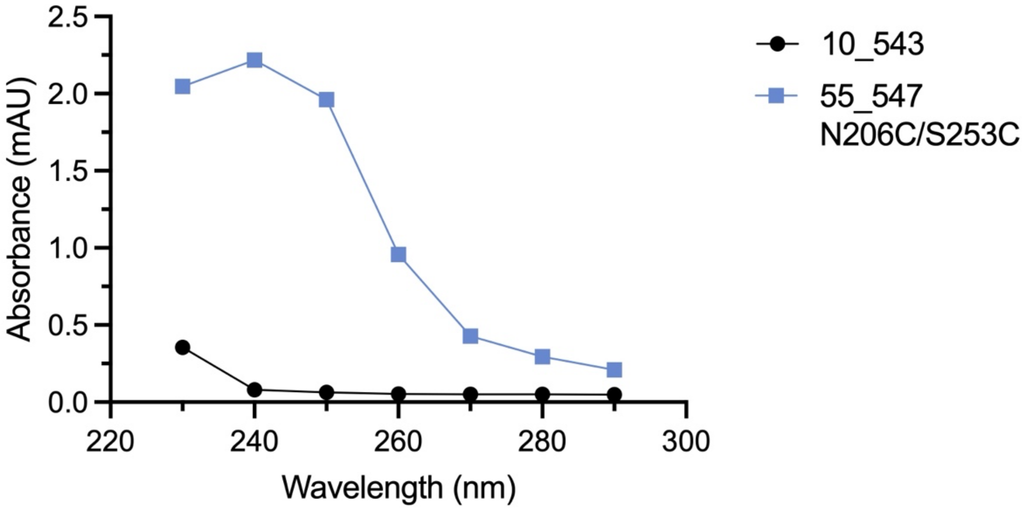


**Supplementary Figure 2.** UV absorbance traces of representative inactive (10_543) and active (55_547 N206C/S253C) between 230 - 290 nm. Absorbance at 260 nm was used for comparison of all variants.

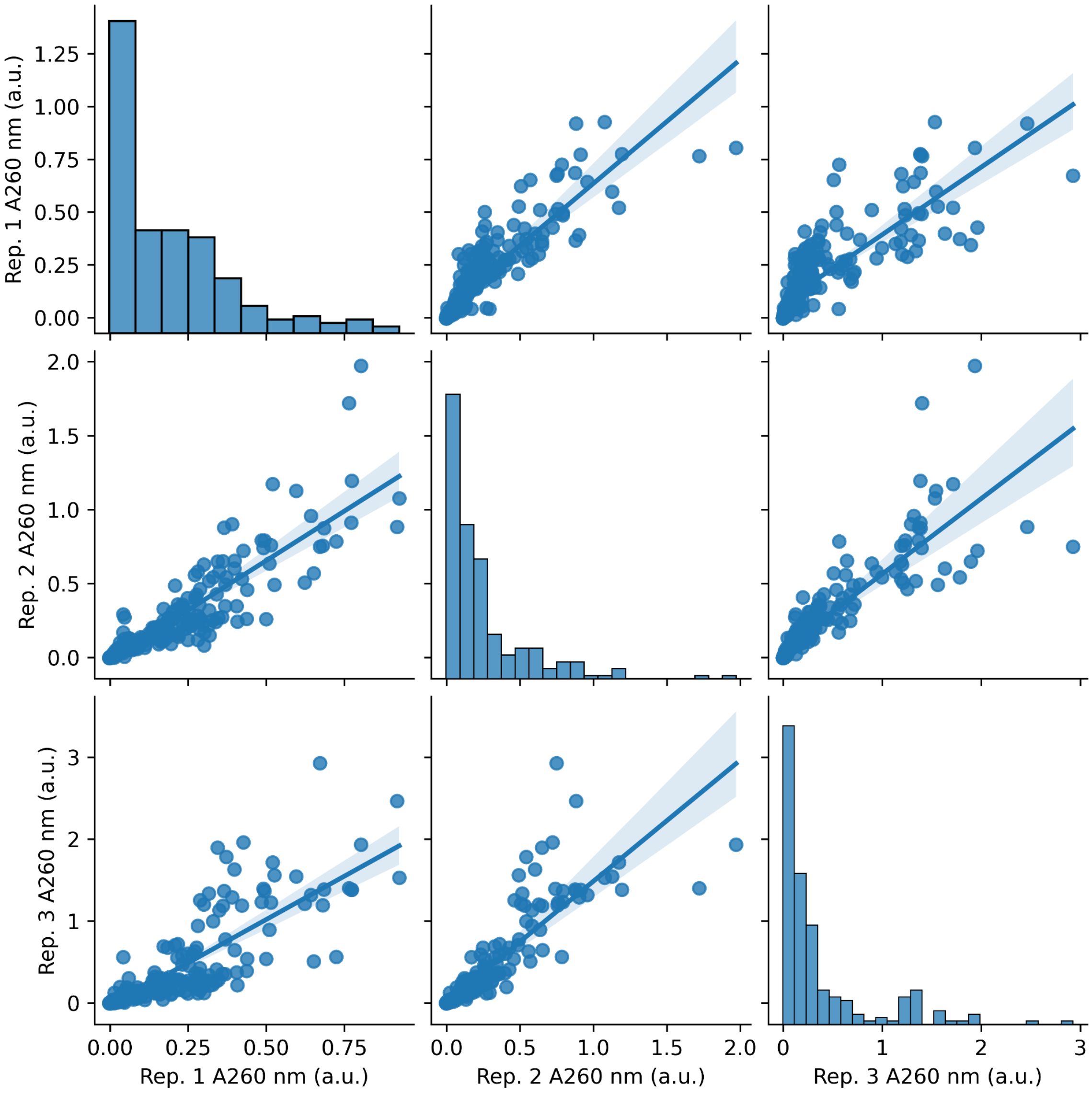


**Supplementary Figure 3.** Correlogram of replicates prior to scaling. All replicates are significantly correlated prior to normalisation by scaling by a constant.

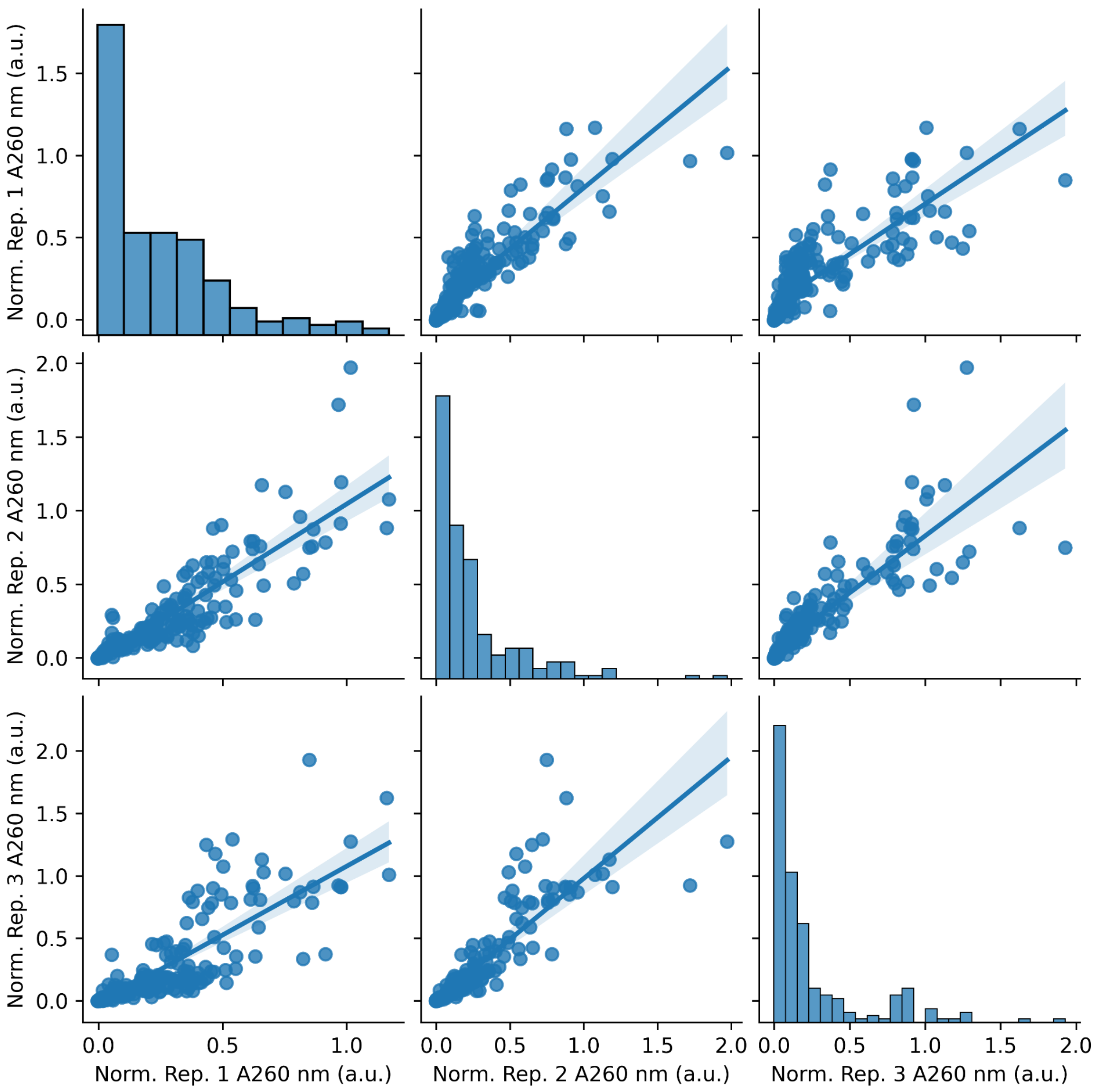


**Supplementary Figure 4.** Correlogram of replicates post scaling. During scaling, replicate 1 and 3 were scaled such that the mean absorbance value of each equaled the mean absorbance of replicate 2 (not scaled). The monotonic (rank) correlation between replicates remains constant with scaling, however, the magnitude of absolute values is reduced by a constant factor (relative to the non-normalised measurements).

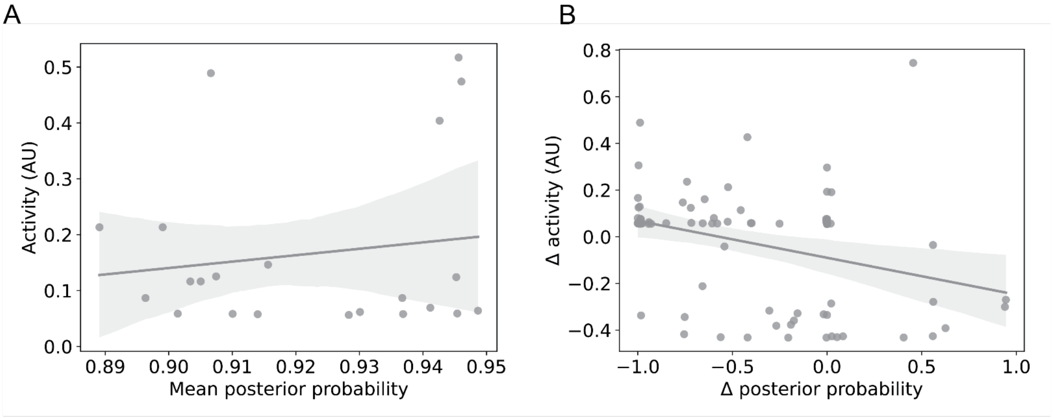


**Supplementary Figure 5.** Posterior probability plotted against PETase activity. **A.** Mean posterior probability plotted against PETase activity over the full dataset of MAP ancestral sequences. **B.** Relative difference in posterior probability plotted against the relative difference in activity from ambiguous alternate ancestral sequences in the backgrounds of 35_442 and 55_547. Posterior probability is a poor indicator of PETase activity.

**Supplementary table 1.** Crystallographic conditions and data. Crystallisation statistical values for the ancestral variants with the highest resolution shell data displayed in brackets.

| | **55_547** | **35_442** |
|---|---|---|
| **Crystallisation Conditions** | | |
| Salt | 0.2 M sodium/ Potassium tartrate | 0.2 M sodium malonate |
| Buffer | 0.1 M BisTris propane pH 7.5 | 0.1 M BisTris propane pH 6.5 |
| Precipitant | 20% (w/v) PEG 3350 | 20% (w/v) PEG 3350 |
| Protein Concentration (mg/mL) | 15 | 30.7 |
| **Data Collection** | | |
| **PDB ID** | **8ETX** | **8ETY** |
| Wavelength (nm) | 0.95373 | 0.95373 |
| Space group | C 2 2 2 | C 2 2 2 |
| Cell dimensions | | |
| *a, b, c* (Å) | 70.2, 150.6, 55.7 | 69.8, 148.5, 55.6 |
| α, β, γ (°) | 90.0, 90.0, 90.0 | 90.0, 90.0, 90.0 |
| Resolution (*Å*) | 41.90 – 1.75 (1.78 – 1.75) | 40.37 – 1.54 (1.57 – 1.54) |
| $R_{merge}$ | 0.159 (2.422) | 0.188 (5.988) |
| $R_{pim}$ | 0.053 (0.795) | 0.054 (1.715) |
| I/σI | 9.1 (1.0) | 10.6 (1.0) |
| $CC_{1/2}$ | 0.997 (0.442) | 0.998 (0.385) |
| Completeness (%) | 100.0 (100.0) | 100.0 (100.0) |
| Redundancy | 9.9 (10.2) | 13.0 (13.4) |
| **Refinement** | | |
| Resolution (*Å*) | 41.90 – 1.75 (1.81 – 1.75) | 40.37 – 1.54 (1.60 – 1.54) |
| No. reflections | 30209 (2970) | 43071 (4210) |
| Rwork/Rfree | 0.1658/0.1937 (0.2907/0.3171) | 0.1694/0.1991 (0.2744/0.2906) |
| No. atoms | 2312 | 2469 |
| Protein | 2000 | 2073 |
| Ligand/ion | 64 | 74 |
| Water | 278 | 322 |

| | | |
|---|---|---|
| B-factors (overall) | 27.41 | 26.15 |
| Protein | 26.09 | 23.50 |
| Ligand/ion | 44.19 | 47.46 |
| Water | 34.85 | 38.34 |
| RMSD | | |
| Bond lengths (Å) | 0.001 | 0.007 |
| Bond angles (°) | 0.579 | 0.877 |
| Ramachandran plot | | |
| Preferred (%) | 98.46 | 98.46 |
| Allowed (%) | 1.54 | 1.54 |
| Outliers (%) | 0.00 | 0.00 |

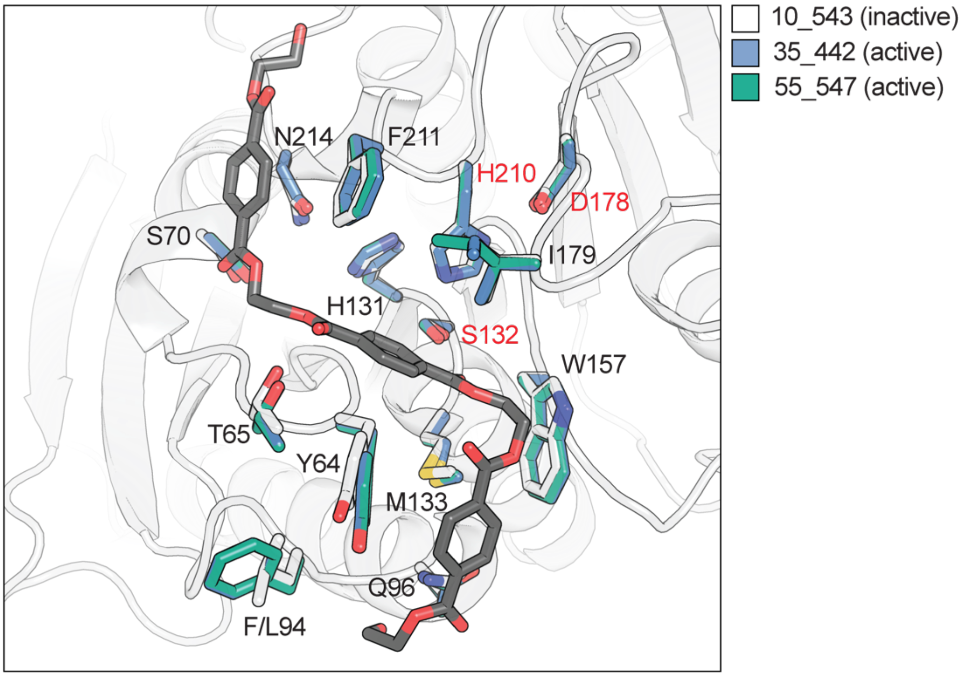


**Supplementary Figure 6**. Comparison of active site and binding site between AlphaFold models of the inactive ancestor 10_543 (white), and active ancestors, 35_442 (blue) and 55_547 (green). Structural analysis reveals uniform alignment of all active site residues (red labels; H210, S132, D178), indicating that PETase activity in the ancestral cutinases does not arise by modifications within the active sites. The docked pose of 2HE-$(MHET)_3$ is shown in a close-up of the binding and catalytic site, and residues that form the putative PET binding site based on homology to LCC are shown.

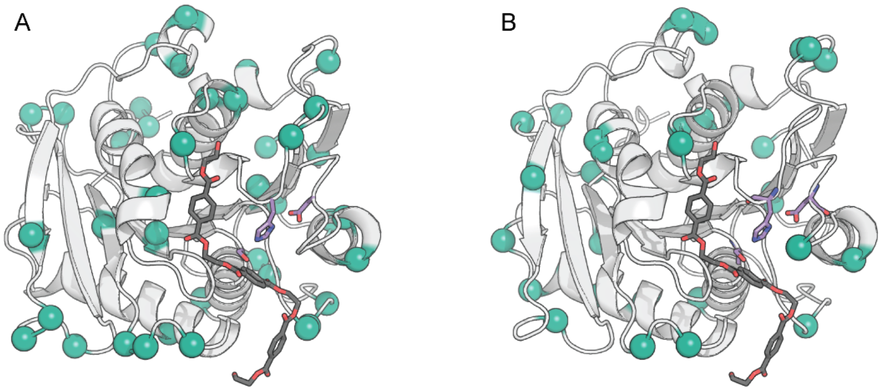


**Supplementary Figure 7.** Ambiguously reconstructed positions in ancestor (**A**) 55_547 and (**B**) 35_442. Positions are highlighted as teal spheres and the docked pose of 2HE-(MHET)$_3$ is shown. Ambiguously reconstructed residues sampled for experimental characterisation are distributed across the protein structure.

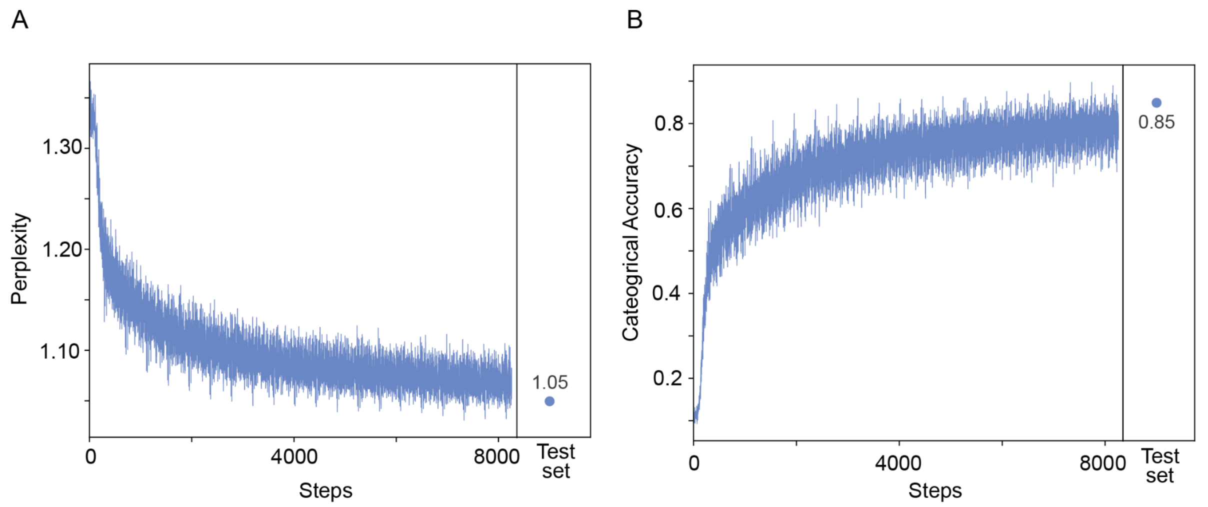


**Supplementary Figure 8. Performance of LASE model over training.** Train and test (10% of available data) (**A**) perplexity and (**B**) categorical accuracy of LASE model after training for 100 epochs. The LASE model (3-layer transformer with 2-head attention and a hidden dimension of 64) was trained with the Adam optimizer and categorical cross-entropy as the loss. Both perplexity and accuracy represent the performance on the masked tokens only (a random 15% of the amino acids from each sequence).

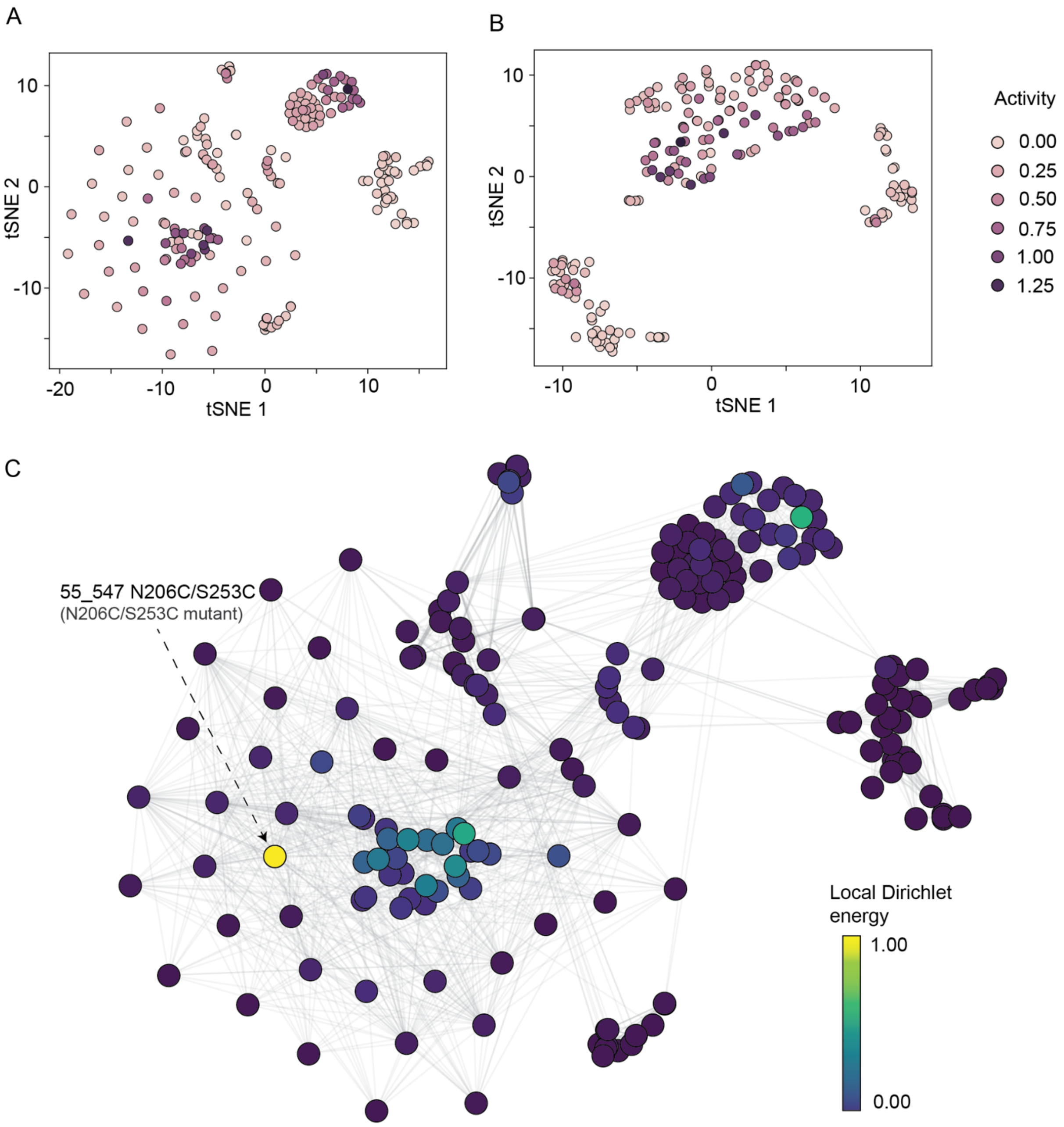


**Supplementary Figure 9.** OHE landscape analysis of PET competent cutinases. Sequence fitnesses shown in (**A**) OHE basis and (**B**) LASE representation basis. (**C**) Local Dirichlet energy shown per node in a OHE encoding basis. Ancestor 55_547_E28Q/E14D/S196A is both the highest energy node and the most fit variant in the OHE representation space.
